# SynBio in 3D: the first synthetic genetic circuit as a 3D-printed STEM educational resource

**DOI:** 10.1101/2022.06.06.493117

**Authors:** Heloísa Oss Boll, Matheus de Castro Leitão, Aisel Valle Garay, Cíntia Marques Coelho

## Abstract

Synthetic biology is a new area of science that operates at the intersection of engineering and biology and aims to design and synthesize living organisms and systems to perform new or improved functions. This novel biological area is considered of extreme relevance for the development of solutions to global problems. However, its teaching is often inaccessible to students, since many educational resources and methodological procedures are not available for understanding of complex molecular processes. On the other hand, digital fabrication tools, which allow the creation of 3D objects, are increasingly used for educational purposes, and several computational structures of molecular components commonly used in synthetic biology processes are deposited in open databases. Therefore, we hypothesize that the creation of biomolecular structures models by handling 3D physical objects using computer-assisted design (CAD) and additive fabrication based on 3D printing could help professors in synthetic biology teaching. In this sense, the present work describes the design and 3D print of the molecular models of the first synthetic genetic circuit, the toggle switch, which can be freely downloaded and used by teachers to facilitate the training of STEM students in synthetic biology.

## Introduction: Background and rationale for the educational activity innovation

Synthetic biology comprises a series of disruptive technologies capable of providing new solutions to global challenges in health, agriculture, industry, and environment (Cameron et al., 2014; Bueso and Tangney, 2017; French, 2019). In its top-down approaches, such area uses molecular biology tools and techniques coupled with standardized engineering principles to design and characterize biological parts, synthetic biological circuits, and systems. Thus, synthetic biology can assign new functions to organisms and redesign pre-existing biological systems to improve features of interest (Fu, 2006; Huang et al., 2010).

Jacob and Monod described, in the 1960s, the first genetic circuit, as they observed that inducible systems, such as the lactose operon from *Escherichia coli*, and repressible systems, like the tryptophan operon or lysogenic systems, respond to similar controlling elements, organized in different genetic circuits (Jacob and Monod, 1961). For example, in the lactose operon, the authors observed that the lactose catabolism was triggered only by a specific inducer molecule, structurally similar to lactose, which stimulated the coordinated synthesis of enzymes that allow lactose processing. On the other hand, when lactose is absent and glucose is present in the medium, the last sugar is preferred, due to easier processing, and the lac operon is silenced by the association of a repressor protein with the operator region of the lac promoter. The description of inducer and repressor components that could genetically control cellular machineries subsequently enabled, decades later, the development of synthetic biological circuits.

In the year 2000, researchers assembled the first synthetic genetic switch, based on two genes that repress each other (Gardner et al., 2000). The application of a specific stimulus induces the expression of one of the genes, and its genetic product inhibits the expression of the other, and vice-versa. One of the genes was placed *in tandem* with a reporter sequence encoding the green fluorescent protein (GFP), and thus the two possible states of the system were the presence or absence of green fluorescence. Due to the similarity with an electronic toggle switch, which can have two possible states - on (light) or off (no light) - their system is also recognized as the genetic toggle switch. Shortly after, Elowitz and Leibler developed the genetic oscillator, a circuit characterized by the interactions among three different genetic repressors, one of which also regulated GFP expression (Elowitz and Leibler, 2000). At a cellular level, the circuit caused periodic oscillations in the detection of green fluorescence, similar to what happens in electronic oscillators. Both studies strengthened the foundations of synthetic biology and demonstrated that it is possible to apply engineering principles to create synthetic biological circuits in living organisms.

Although synthetic biology has many similarities with engineering, the same is not observed when concerning education (Diep et al., 2020). Electrical engineering students, for example, can usually obtain parts of electrical circuit, such as conductor wires, electronic switches, and LEDs, to learn with practice the concepts studied during theoretical classes - such as building a simple electronic toggle switch to turn on a light bulb (Kansal creation, 2019). In contrast, synthetic biology focuses on designing and building biological circuits which operate at the molecular scale. Consequently, it’s harder for students to connect the concepts learned in class with the actual dimension of subcellular processes due to the scarcity of specific pedagogical methods.

In this way, the teaching of synthetic biology faces challenges quite similar to those outlined for molecular sciences education. For example, the complex process of transforming two-dimensional, static drawings present in books and scientific articles into three-dimensional, dynamic models (Wu et al., 2001). This skill is implicitly requested throughout the life sciences academic curriculum, and the fact that a substantial number of students have difficulties with three-dimensional mental visualizations is often neglected, which can bring prejudice to their professional careers in Science, Technology, Engineering and Mathematics (STEM) fields (Pittalis and Christou, 2010). In addition, most methods proposed for teaching synthetic biology until then are using computer programs that allow the visualization of molecular reactions or performing molecular experiments in the laboratory. BioBits, for example, is a modular kit developed for synthetic biology education that sensorially engages students through experiments that produce fluorescence, fragrances, and hydrogels (Huang et al., 2018). However, it does not allow the visualization of the molecular processes as they occur, nor the molecular structures involved.

Nevertheless, since the late 40s, Robert Core and Linus Pauling, pioneers in studies of protein structures, developed the first representation of macromolecules in terms of complex space-filling models (Paulin and Corey, 1950; Paulin, 2015). This system represents atoms of different chemical elements as spheres of different colors whose diameter is proportional to their atomic radius (Walter, 1965; Olson, 2018). In 1957 researchers reported the first experimental structure of a macromolecule (Kendrew et al., 1958), and, for twenty years, physical models were the principal tool for representing the structures of biological macromolecules (Olson, 2018). Physical models fell into disuse during the early 1980s, when the visualization of biomolecular structures by molecular computer graphics software became popular. Fortunately, more recently, several independent efforts created physical models of proteins using new rapid prototyping technologies based on 3D printing. 3D printers enable the manufacture of macromolecules structures using digital atomic coordinates as a .pdb file downloaded from the Protein Data Bank (PDB), with low waste and high precision and efficiency, simplifying the process of fabricating molecular models.

Considering education, according to Salzman et al. (1999), students who struggled to understand molecular biological concepts perceived more valuable tactile feedback than isolated 3D representations. Furthermore, the use of three-dimensional educational materials is particularly significant to visually impaired students, who face substantial barriers in the classroom due to the lack of tactile methods, which are fundamental for a better understanding of biological concepts (Stone et al., 2020). In this way, the present project aimed to facilitate synthetic biology teaching through computational manipulation and three-dimensional printing of biological structures involved in the first synthetic biological circuit, the genetic toggle switch.

With this resource, professors would be able to use parts on an adequate scale allowing students to manipulate and understand the theoretical concepts studied in the classroom in a practical way. By using 3D objects, students would be able to understand the three-dimensional properties and the relative size of molecules. These tools could help professors and students to improve their comprehension of complex concepts in structural chemistry and biology. The physical models would provide a tactile and visual input sensing combination for teaching and understanding macromolecules spatial relationships and mechanisms in ways that illustrations and computational models cannot.

## Pedagogical framework(s), pedagogical principles, competencies/standards underlying the educational activity

Our methodology is focused on providing professors with a detailed description of a tactile-visual strategy for teaching synthetic biological circuits. The approach proposed here has already been tested by other authors in the broader field of STEM education, which includes synthetic biology. Some significant findings highlighted in previous studies are that (1) active engagement learning strategies have already been shown to reduce the percentage of failure rates compared to regular lectures in undergraduate courses across STEM disciplines; (2) it is associated with a statistically significant improvement in individual learning performance and an increase in the average assessment scores in Molecular Sciences (Newman et al., 2018). Specifically in Molecular Biology, the use of physical models has been shown to increase learning gains related to the central dogma of molecular biology, and we hypothesize that this is also true for synthetic biology due to the fundamental similarities between the areas (Gordy, 2020).

For the construction of our educational resource, we elected the first genetic toggle switch, designed, synthesized, and successfully tested by Gardner and collaborators (Gardner et al., 2000). Foremost, we identified the molecular elements that compose this genetic circuit, two molecular inducers, aTc (anhydrotetracycline) and IPTG (isopropyl β-D-1-thiogalactopyranoside); two repressor gene products, LacI (lactose operon repressor) and TetR (Tet repressor protein); and a reporter gene product, GFP (green fluorescent protein). As the system depends on gene expression, we also included an RNA polymerase to allow the understanding of the transcriptional activity control and generic DNA strands composed of four modules each to represent the two mutually repressive genetic cassettes.

Figure 1 shows a schematic representation of the steps to obtain the 3D-printed molecules of the first genetic toggle switch. Digital 3D atomic coordinates as a .*pdb* file were downloaded from Protein Data Bank (PDB) (http://www.pdb.org/pdb/), repeating the process in the Molview database and downloading the files in *Mol* format. Structures were prepared in the PyMol and ChimeraX v1.3 (https://pymol.org/2/, https://www.rbvi.ucsf.edu/chimerax) software by extracting unitary cell, to remove waters, ligands, and calculate the molecular surfaces. Afterwards, molecular surfaces structures were exported as .*stl* file using ChimeraX or as VRML 2 file from PyMol to repair and hollowed with Autodesk Meshmixer v.3.5.474 or Blender software. In Blender, each element was imported individually as X3D Extensible 3D files. The structural skeleton of each piece was deleted, preserving only the empty surface shells. In Meshmixer the molecular surfaces .*stl* file were hollowed and repaired to generate a new .*stl* file for use in UltimakerCura slicer.

**Figure 1.**
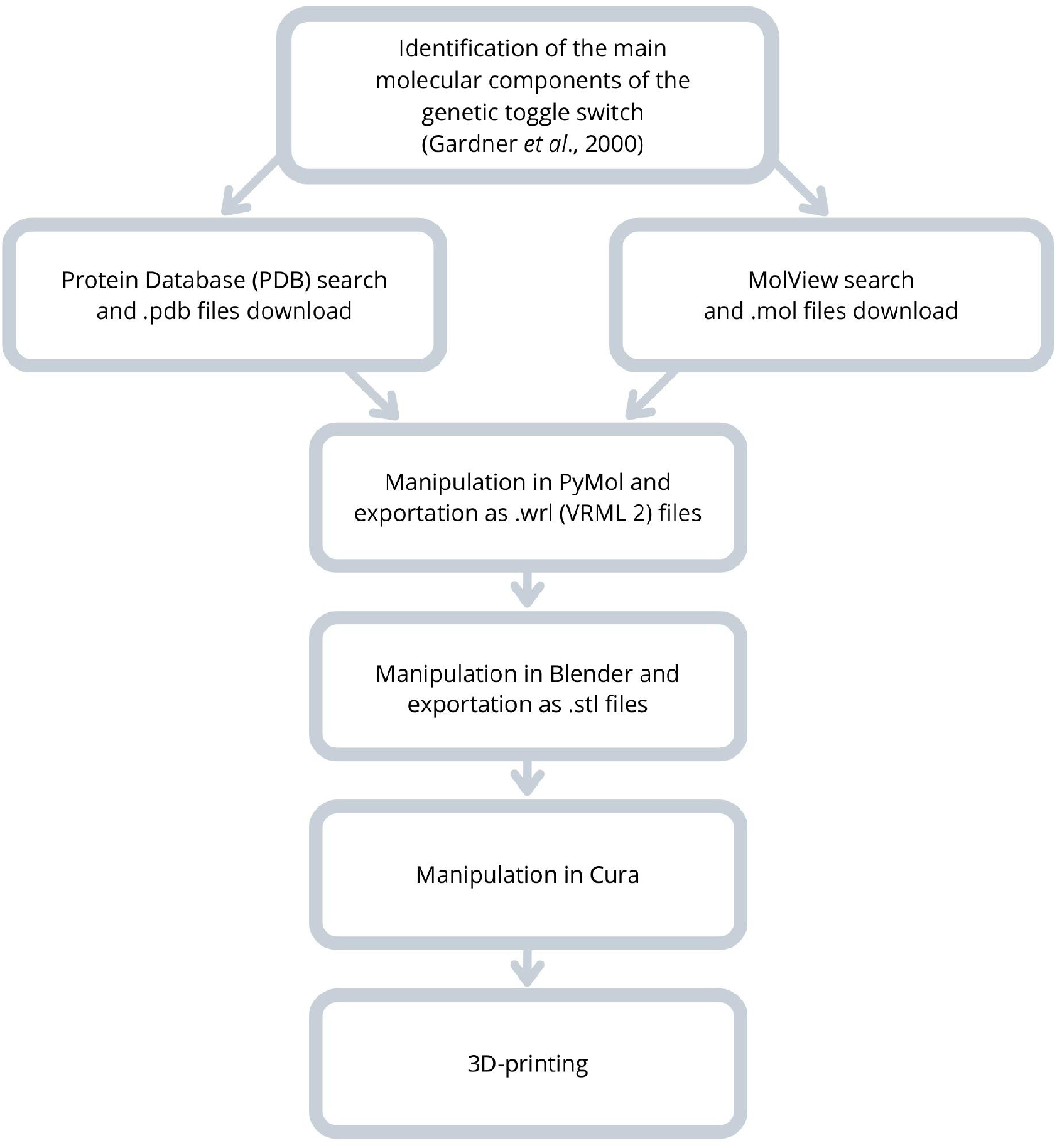
Schematic representation of the pipeline used for production of the genetic toggle switch as a system of 3D-printed pieces.

During the slicing onto UltimakerCura software, we scaled the macromolecules from .*stl* files into two size scales. Inducers were printed at 100% of the initial size and the other components at 70% of the scale. The 3D prints were performed on a fused deposition modeling (FDM) 3D printer with an extrusion nozzle of 0.4 mm in diameter using plastic polylactic acid (PLA) filament of 1.75 mm were used. In the slicing software, we set the following parameters (1) the extrusion width was adjusted according to the extrusion nozzle diameter; (2) the printing temperatures ranged from 200 to 215 °C, (3) the printing speed ranged from 20 to 60 mm/s, with the temperature and speed parameters changed in direct relation to the individual size of each printed part. We used 200g of PLA filaments in yellow (lacI), white (IPTG), blue (TetR), light blue (aTc), black (DNA) and green (GFP). For subunits of RNA polymerase different colors were used in a total mass of 543g. After printing, print supports were removed by hand from the models and small 5×5 mm neodymium magnets were added to allow the reversible connection between the pieces representing the molecules of the toggle switch and better simulate the behaviors of transcription activation and repression. Subunits of RNA polymerase were assembled with glue for plastic.

## Learning environment (setting, students, faculty); learning objectives; pedagogical format

After 3D-printing the files, we developed a series of steps that the responsible teacher should go through to explain how the synthetic circuit works in a simple, straightforward way. First, we recommend dividing the students into small teams, with a maximum of 4 participants. Then, the teacher will reproduce the following steps in front of the class:

1. Introduce all the molecules and genetic parts involved in the synthetic genetic circuit, describing that IPTG and aTc will be the inducers and LacI and TetR will be the repressor proteins for pTrc-2 and P_L_tetO-1, respectively. As promoters, P_L_tetO-1 and pTrc-2 control the expression of LacI and TetR together with GFP, respectively, in the two different expression cassettes.
2. Simulate how RNA polymerase transcribes both of the cassettes. In the first case, the magnetic connection of the RNA polymerase with the pTrc-2 promoter region of a DNA strand causes the expression of TetR, a repressor protein, and GFP, a green fluorescent protein. In the second case, the transcription leads to the expression of LacI, another repressor. These final products should not be visible to the class until transcription has been simulated. However, the translation process omitted in our system should be highlighted as the step responsible for converting the information contained in the expressed mRNAs into individual proteins.
3. Show how the addition of the two repressors blocks RNA polymerase activity at each of the two promoters in the two expression cassettes. Repeat the last step, but explain that a physical attachment of LacI or TetR on the promoter region of pTrc-2 or P_L_tetO-1, respectively, impedes the transcription of each cassette by RNA polymerase. No transcription and consequently no translation takes place, so the class does not get to see the final product. It is important to note that in the system, the binding of only one molecular repressor per promoter is a simplified representation of the cell’s reality; many more molecules operate to block promoter regions.
4. Explain how the two cassettes interact with each other.
  a. Draw a parallel with a standard light switch that can set a simple electrical system into two possible states: the presence (ON) or absence (OFF) of light. Thus, the genetic switch can set the biological system into two possible states: the presence (ON) or absence (OFF) of green fluorescence. In this case, it is worth noting that GFP is the product that confers this characteristic.
  b. Draw a parallel between the two cassettes and show that IPTG, once added to the medium, is present inside the cell, and that the repressor product of the second cassette, LacI, has the property of binding to this molecule, using the magnets embedded in both parts. Then, ask the students: if LacI is not bonded to pTrc-2 due to the interaction with IPTG, what happens to the expression of the pTrc-2 cassette? As a consequence, the transcription process must be shown again, which results in the synthesis of TetR and GFP.
  c. Highlight the TetR produced by the first cassette. Explain that this second molecular repressor attaches to the P_L_tetO-1 promoter of the second cassette and blocks its expression. Consequently, LacI is no longer produced. Finally, the teacher should ask: “What state does the system assume when IPTG is added to the medium?” The final answer is: “The system is in its ON state and shows a green fluorescence”.
  d. Explain how the second state is reached by showing the property of the molecular repressor aTc to bind to TetR by a magnetic connection. Then the teacher should ask the students: “If, after its addition to the medium, aTc interacts with TetR, what happens to the expression of the P_L_tetO-1 cassette?”. In this case, the transcription process is restarted, leading to the synthesis of LacI, a repressor of the pTrc-2 promoter. When the pTrc-2 is blocked, there is no expression of GFP and TetR. Finally, the teacher should ask: “What state does the system assume when aTc is present in the medium?”. The final answer is: “The system is OFF and colorless”.

After the main explanation by the teacher, each small group should be allowed to manipulate the 3D printed pieces. We also suggest that the teacher concedes each group a few minutes to try to reproduce what they have learned and check if any concept was not fully understood. Students could then alternate and give a short presentation to the whole class, explaining how the different molecules control the gene expression on the synthetic genetic circuit.

## Results to date/assessment (processes and tools; data planned or already gathered)

Although Jacob and Monod had pointed out in the ‘60s the existence of biological regulatory circuits, it was not until the beginning of the new millenium that Gardner and collaborators reported the first design and test of a synthetic genetic circuit, the toggle switch (Gardner et al., 2000).

The rationale for this proposal relies on the hypothesis that professors will benefit from 3D models to better explain concepts of genetic circuits. We selected the toggle switch as a starting point, not only because it was the first synthetic circuit assembled, which paved the way for further designs, but also due to its simplicity: it involves only a negative control that relies on two repressors to regulate the expression of the GFP gene. Figure 2 describes all genetic parts that comprise this switch, comparing the computational models experimentally obtained and submitted online and the 3D-printed pieces. By preserving the dimensions of each molecule, molecular inducers (IPTG and aTc) are significantly smaller than protein repressors (TetR and LacI). Also, we used filaments colored in similar tones to the products derived from each of the cassettes, so students can easily associate which system is which.

**Figure 2.**
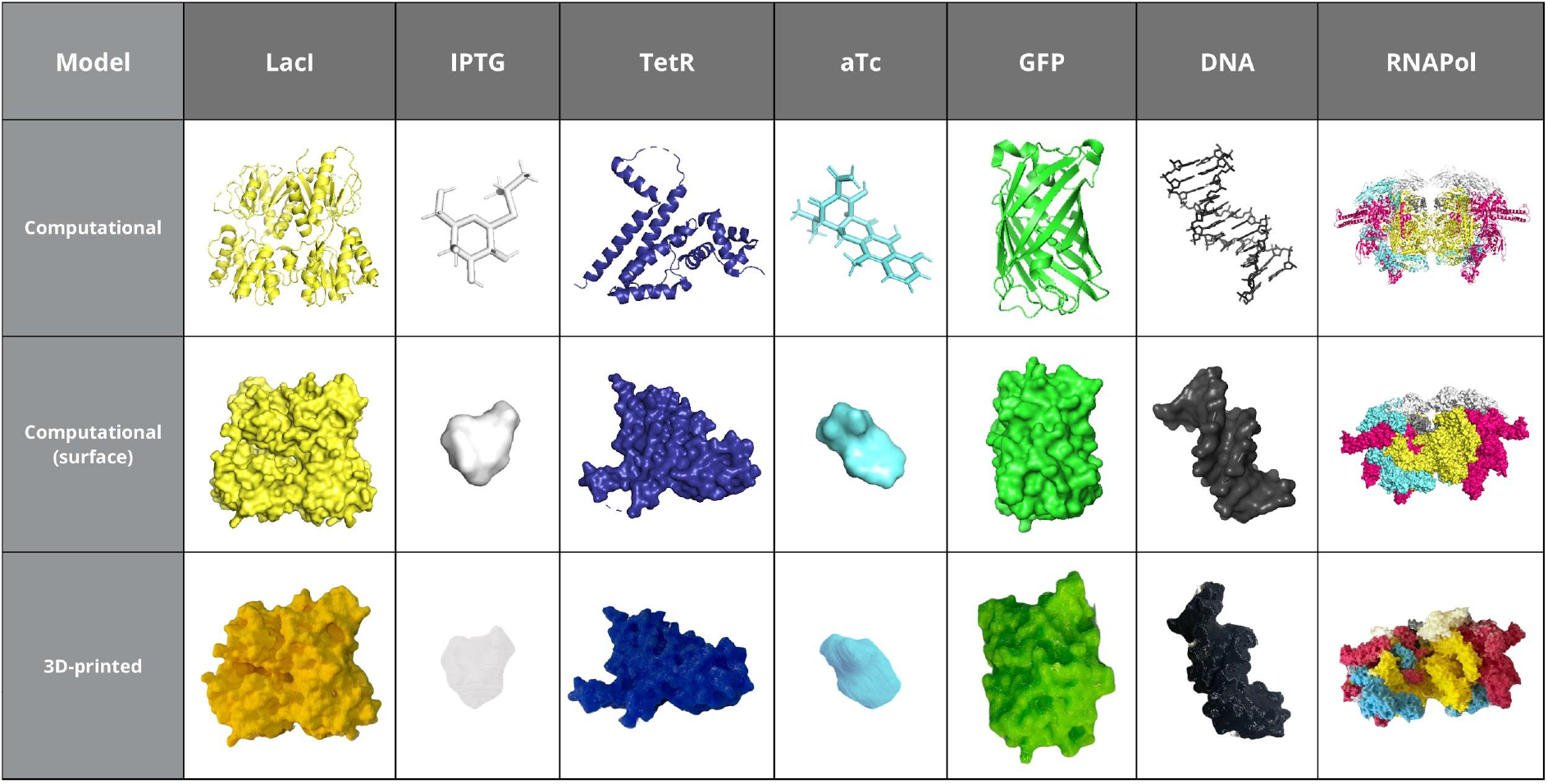
Visual comparison of 3D printed models that compose the genetic toggle switch in relation to their respective computational structures.

After preparation, the final genetic circuit elements were 3D-printed and finalized with magnets. All files are available in the Supplementary Material, on our laboratory’s website, synbiolabunb.com, and under the CC BY 4.0 license. This authorizes the work to be shared and adapted under conditions of giving credit to the original authors, not using the materials for commercial purposes, and not applying legal terms or technological measures that legally restrict others from doing anything the license permits.

Figure 3 describes the different cellular states dependent on its intracellular inducer content, showing how the genetic control expression is linked to Boolean algebra function as binary variables, considering 1 when the gene is expressed and 0 when it is not. First, it explains the design of the synthetic genetic circuit to control the expression and accumulation of the reporter protein, GFP. With the IPTG addition to the medium, it diffuses within the cells and binds to the LacI protein. This way, the operator of the pTrc-2 promoter is free, allowing the flow of the RNA polymerase - a process called a transcription “current”, analogous to an electric current as a transistor-like device, named transcriptor. The result is the expression of the cassette composed of TetR and GFP. TetR binds to the promoter of the other cassette, P_L_tetO-1, blocking the expression of LacI, and GFP sets the “ON” state of the system, making the cells turn fluorescent green. Then this figure illustrates how the cells can change state and turn colorless. When aTc is added to the medium, it enters the cell and binds to the TetR protein, leaving the operator of the P_L_tetO-1 promoter available for the RNA polymerase to transcribe. Hence, LacI is produced, and it blocks the promoter of the other cassette, pTrc-2, setting the “OFF” state of the system: that is, cells are unable to express GFP and turn colorless. In summary: IPTG causes the cells to turn fluorescent green, whereas aTc has the opposite effect, maintaining them colorless.

**Figure 3.**
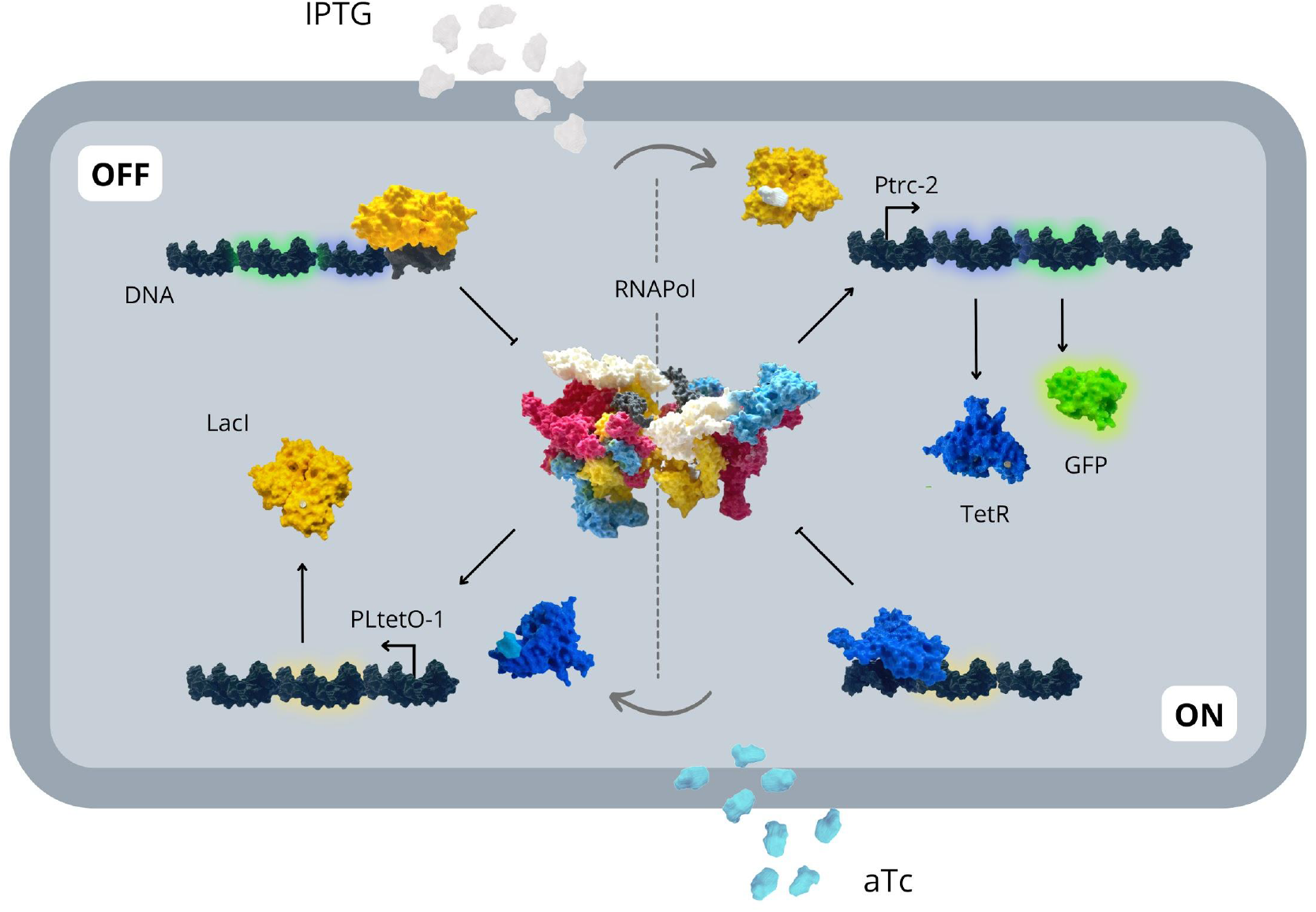
Different cellular states that compose the toggle genetic circuit in 3D printed molecules. The molecules and the genetic parts needed in the synthetic genetic circuit to turn it on its ON state are: IPTG (the inducer), LacI (repressor protein) and pTrc-2 (promoter). In its initial state, the LacI repressor is bound to the pTrc-2 promoter, blocking the RNA polimerase flow. After adding IPTG to the medium, it diffuses into the cellular environment, binds the LacI protein, releasing it from the pTrc-2 promoter, and allowing the function of the transcriptor. The cell is now in its ON state, expressing GFP and TetR. The TetR repressor protein binds to the P_L_tetO-1 promoter and blocks the RNA polimerase flow along the cassette. Then, now, to switch to the OFF state the molecules and the genetic parts needed will be aTc (inducer), TetR (repressor protein) and P_L_tetO-1 (promoter). Once the aTc is added to the medium, it diffuses into the cellular environment, binds the TetR protein, releasing it from the P_L_tetO-1 promoter, and allowing the function of the transcriptor. The cell is now in its OFF state, expressing LacI and thus repressing the first cassette from transcribing and traducing TetR and GFP.

The 3D printed biological molecules that compose the first synthetic genetic circuit were presented to and evaluated by undergraduate students attending the Genetic Engineering class at the University of Brasilia, following the series of steps described above that the responsible professor must follow to explain how the synthetic circuit works. The results showed that approximately 46% of the students found it difficult or very difficult to understand the switch before the class, and 38.5% had an average understanding (Supplementary Figure 1A). Almost 80% stated that it was not easy to understand the results of the first synthetic gene switch relying only on the explanatory figures in the articles (Supplementary Figure 1B). Students also reported that the 3D printed biological molecules helped them very much in understanding the switch. Only 1 student stated that the printed resource was not helpful; but this student also reported that the class dynamics were good (Supplementary Figure 1C). In addition, approximately 90% of them stated that they strongly believe that using 3D printed molecular structures would make it easier to understand other systems related to the field of genetic engineering (Supplementary Figure 1D).

## Discussion on the practical implications, objectives and lessons learned

Genetic circuits are a combinatorial network of switches that can perceive different inputs, process the information, and generate outputs. Since the beginning of the 2000s, several simple synthetic circuits have been designed, built, and tested (Gardner et al., 2000; Elowitz and Leibler, 2000). These were the cornerstone for assembling more complex ones, which led to further implications in biotechnology, medicine, agriculture, and food sectors (Khalil and Collin, 2010), bringing the synthetic biology area to a pivotal importance in the 21st century (Karoui et al., 2019). Therefore, it is of great interest that future researchers in the different areas of science have a profound understanding of genetic circuits.

The traditional educational system still fosters a passive learning process in which students are stimulated to hear explanations, take notes and then take exams to prove their knowledge. However, we hypothesized that complex and emerging areas, such as synthetic biology, cannot be fully understood without a modern and practical educational approach. The process of elaborating a synthetic biology project is deeply amalgamated into digital technologies, such as apps made to design plasmids from scratch and machines that automate strain engineering. Hence, classes dedicated to synthetic biology education also need to adapt and integrate the use of technology, following up with the tendency and allowing students to usufruct from active learning methodologies. Moreover, once the 3D-printed molecular structures from the first synthetic genetic circuit was presented to undergraduate students approximately 80% of them reported that this educational resource helped them on the better understanding of the toggle switch.

In conclusion, the method showcased in this article has as a foundation the combination of digital manufacturing processes, which can be used to make (almost) anything, anywhere, with readily-available molecular computational elements (Gershenfeld, 2012). Digital fabrication is a design and production process focused on turning bits into atoms, increasingly present in academic and school settings. On the other hand, databases like PDB are free to use and reunite thousands of digital molecular components, which allows the exportation in formats compatible with 3D printers. Bridging the gap on how simple the process of connecting those two resources is a crucial step to incentivizing teachers to explore it and also develop their systems for their classes.

## Acknowledgment of any conceptual, methodological, environmental, or material constraints

We had some technical issues during the 3D-printing testing phase due to 1) .*stl* files were too large and did not load in the Ultimaker Cura; 2) the settings of the printer weren’t ideal, and the resulting pieces were low in quality. We solved these issues by reducing the size of the files in Blender and exporting it again as a .*stl* file, optimizing the printing supports, and improving the definition of the molecule surface.

## Data availability statements are required for all articles published with Frontiers

Original datasets are publicly available and can be found here: [link/accession number].

## Acknowledgments

We thank our supporting agencies, DPI/DPG (Decanato de Pesquisa e Inovação e Decanato de Pós-Graduação, from the University of Brasilia, Brazil), DEX (Decanato de Extensão, from the University of Brasilia, Brazil), FAPDF (Research Support Foundation of the Federal District, Brazil), and CNPq (National Council for Scientific and Technological), Brazil.We also thank the professors of the Genetic Engineering class, Dr. Ildinete Silva-Pereira, Dr. Marcelo Brígido, and Dr. Juliana Franco for letting us present the 3D-printed toggle switch in their classroom.

## Author contributions

C.M.C conceived the study; H.O.B performed the data collection; H.O.B adapted the data to 3D printer; M.C.L and A.G set up the conditions and printed all the biological structures; H.O.B and C.M.B wrote the manuscript. All the authors analyzed, revised, and discussed the data and contributed to the final paper.

## Competing interests

The authors declare no conflict of interest.

## Notes

https://drive.google.com/drive/folders/1__nhOkkw0pPMu6on_f6zNxE6nZP1mDvW?usp=sharing

### Competing Interest Statement

The authors have declared no competing interest.

