## Supplementary Figure 1 for "SynBio in 3D: the first synthetic genetic circuit as a 3D-printed STEM educational resource"

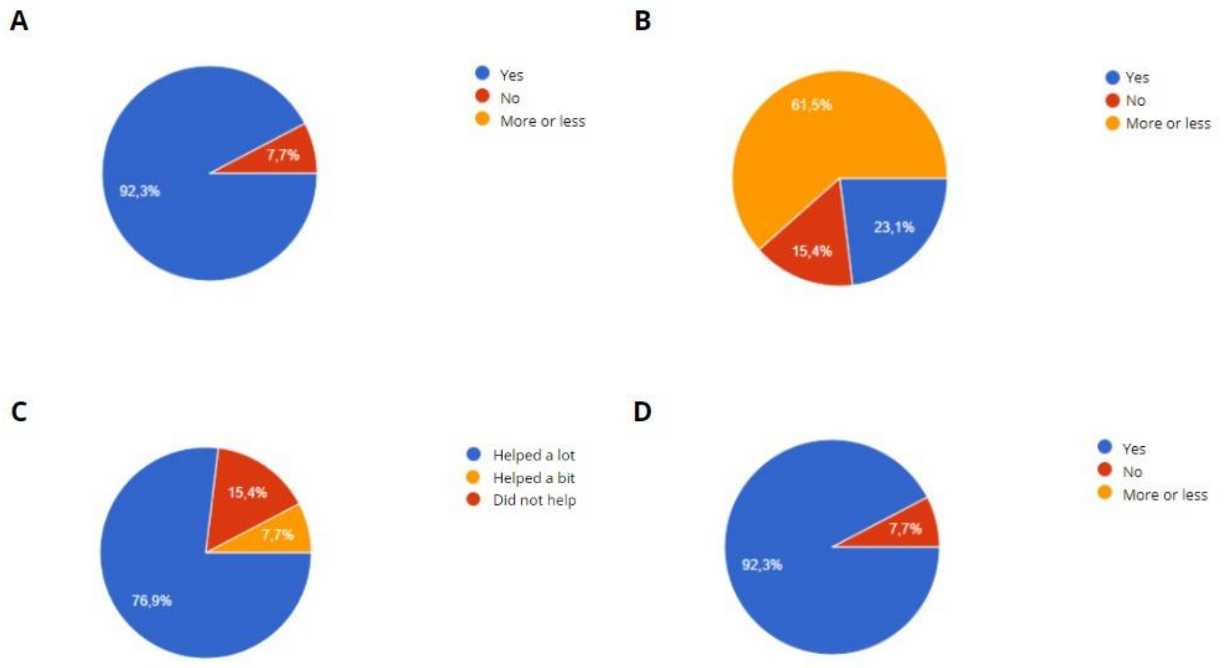


**Supplementary Figure 1**.3D-printed molecular structures that compose the toggle switch presented as a STEM educational resource to undergraduate students from the Genetic Engineering class of the University of Brasília (n=13). (A) Percentage of students reporting their comprehension of the first synthetic genetic circuit before the class that presented the 3D-printed molecular structures of the toggle switch on a scale from 0 (very easy) to 5 (very hard). (B) Percentage of students that reported about the comprehension of the first genetic circuit from the figures presented on the Gardner et al., 2000 article, being blue fully understanding, orange mid-term understanding and red no understanding. (C) Percentage of students reporting their perception about the comprehension of the toggle switch after the 3D-printed molecular structures were presented to them (blue, helped very much; orange, helped; red, did not help). (D) Percentage of students that declared their perception if 3D-printed molecular structures would help the understanding of other topics studied in the Genetic Engineering class, being blue, yes they think it would help them and red they do not think would be of any help.
